# Early-Life Systemic Inflammation Modulate Microglia Phenotype to Slow Down Aβ Pathology in the 5xFAD Mouse Model of Alzheimer’s Disease

**DOI:** 10.1101/2024.11.19.624204

**Authors:** Yiyi Yang, Marta García-Cruzado, Hairuo Zeng, Xinrui Wang, Sara Bachiller, Lluís Camprubí-Ferrer, Bazhena Bahatyrevich-Kharitonik, Rosalía Fernández-Calle, Tomas Deierborg

**Affiliations:** Department of Experimental Medical Science, Experimental Neuroinflammation Laboratory, Lund University; Institute of Biomedicine of Seville (IBiS), Virgen del Rocío University Hospital, University of Seville, CSIC; Department of Medical Biochemistry, Molecular Biology, and Immunology, School of Medicine, University of Seville, Seville, Spain

**Keywords:** LPS, early-infection, inflammation, microglia, monocytes, single-cell sequencing, Alzheimer’s disease

## Abstract

Early infection in life has been implicated in increasing the risk for neurological disorders. Here we performed single-cell sequencing of microglia and monocytes from 6-month-old WT and 5xFAD mice subjected to one dose of LPS (1mg/kg) at postnatal day 9. We successfully mapped disease-associated microglia (DAM) and perivascular macrophages in our data and demonstrated a subpopulation of microglia that adopted a monocyte-like profile, marked by Lyz2, Tmsb10, Lgals1and Lgals3. This unique subset appeared in response to early systemic LPS challenge and AD pathology but diminished in the presence of double stimulus. Different cytokines were altered in the brain and periphery as seen using mesoscale plates. GM-CSF and MIP-1α levels were altered in an amyloid-β(Aβ)-dependent manner in hippocampus. MIP-1β and IFN-γ were altered upon early LPS stimulation. In the periphery, we found MMP-9 was significantly increased in serum samples from 5xFAD mice. Interestingly, early LPS stimulation significantly elevated TNF-α in serum from WT and 5xFAD mice, but was reduced in the hippocampus due to Aβ pathology. The LPS treatment in 5xFAD mice had a tendency to improve the short-term memory deficit. Taken together, we observed long-lasting effects from early life stress, including activation of inflammation in the periphery and brain through modulation of different signaling cascades.

## Introduction

Microglia are the main innate immune cells of the central nervous system (CNS). They continuously survey their microenvironment and have the ability to interact with neurons and regulate their activity^1^. Emerging evidence has shown that microglia are key causative players in neuroinflammation, which in turn is believed to play a major role in neurodegenerative diseases, such as Alzheimer’s disease (AD)^2^. Monocytes from the periphery also play a role as they facilitate communication to the CNS about systemic immunity. However, the contribution of systemic immunity, infiltrating monocytes and microglia to the progression of AD remains controversial ^3–5^.

Epidemiological and experimental data have shown that early exposure to different common pathogens is related to an increased incidence of neurological disorders^6^. It may also have deleterious effects by generating neuroinflammation in the central nervous system (CNS), which may exacerbate the pathology of neurodegenerative diseases^7^. Increasing evidence has demonstrated the heterogeneity of microglia and monocytes under different pathological conditions using single cell RNA sequencing^8–10^. However, the different roles of these two cell types are not well known and not been deciphered due to the difficulty of ambiguous cell markers and common origin of monocytes and microglia.

Thus, our study aims to investigate how systemic inflammation affects the immune system of the CNS and what early alterations occur in microglia and monocytes that may lead to destructive effects in CNS. We also attempt to elucidate the underlying mechanisms and unique profiles of these myeloid cells in wild type (WT) and an AD mouse model (5xFAD).

## Materials and Methods

### Animals

Transgenic 5XFAD mice on a C57/BL6-SJL background were purchased from Jackson Laboratory. These mice express human APP and PSEN1 with three mutations in the human APP transgene (the Swedish mutation, K670N/M671L; the Florida mutation, V716V; and the London mutation, V717I) and two mutations in the human PSEN1 transgene (M146L/L286V). These mutations are related to familial forms of AD, and the transgenes are expressed under the neuron-specific Thy-1 promoter. For this study, male mice were used and analyzed at 6 months of age. Age-matched wild type (WT) littermates were used as controls. Animal housing, handling and experiments were performed under the international guidelines approved by Malmö/Lund animal ethics committee (M30-16, Dnr5.8.18-01107/2018). Mice were housed in groups and maintained in 12 h light/dark cycle with free accessibility to food and water.

### Intraperitoneal injection of LPS

At postnatal day 9, mice were subject to intraperitoneal injection of either lipopolysaccharide (1mg/kg, L4516-from *Escherichia coli* O127:B8, Sigma-Aldrich) or an equal amount of saline for control. All pups were removed from the mother at the same time and returned as a group. All of the mice were induced using the same batch of LPS in this project. No mortality was observed due to this concentration of LPS. The outline of the experiment is shown in Fig.1A. Briefly, mice (n=26) were divided mainly into two groups dependent on their genotypes. To study the effect of early inflammation on monocytes and microglia under AD pathology, the mice were further separated into two additional groups based on LPS treatment (WT-Vehicle, n=6; WT-LPS, n=6; 5xFAD-Vehicle, n=7; 5xFAD-LPS, n=7). At 6 months of age, mice underwent recognition tests described below 48 h before the sacrifice for sample collection.

**Figure 1.**
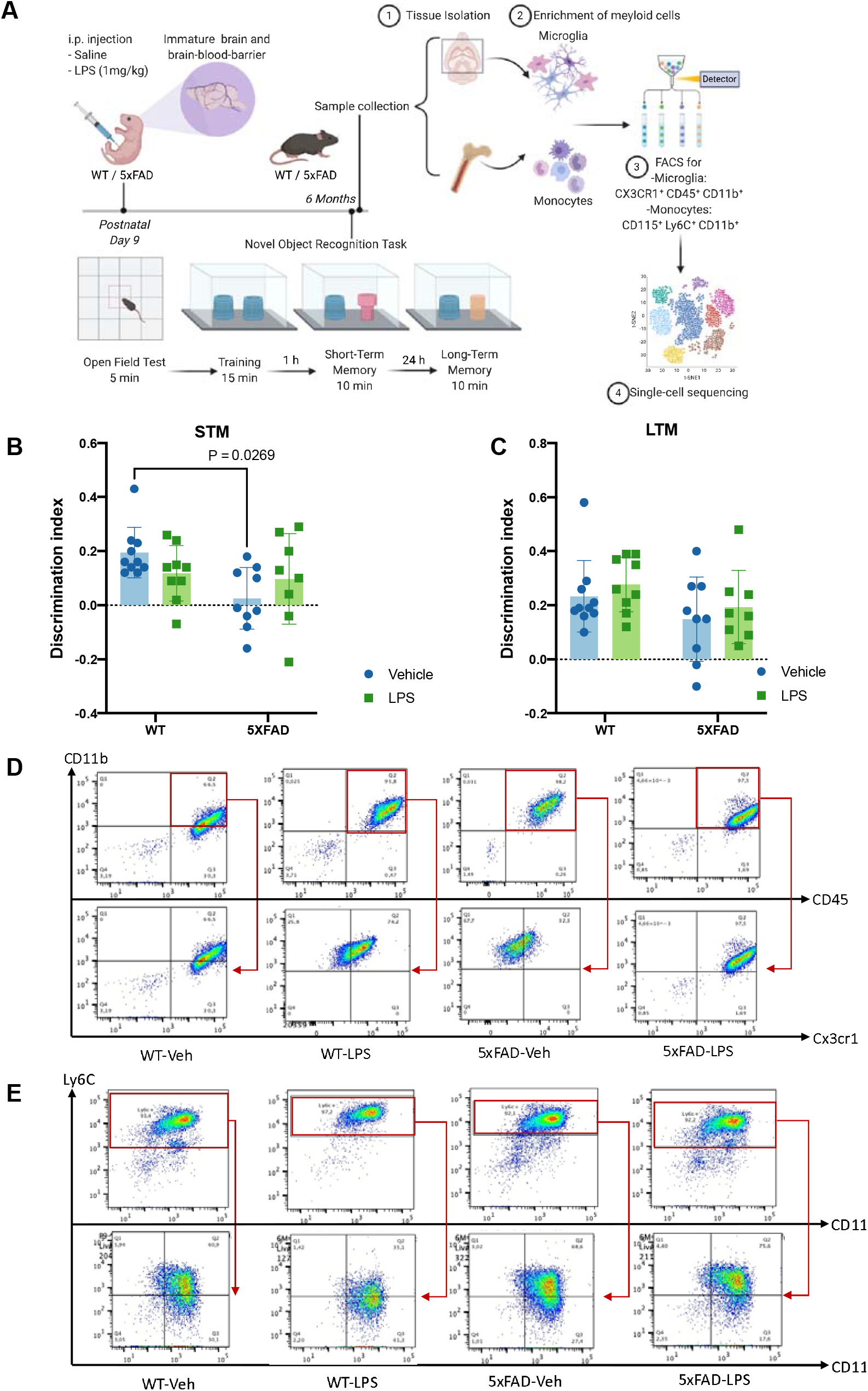
Experimental outline of the study illustrating memory behavioral tests performed in 5xFAD. A) Experimental outline of the study demonstrates the timeline of i.p. injection of LPS (1mg/kg) and procedure of a novel recognition test performed in 6 months old mice. B) Bar graph represents the preference of novel object in four different experimental groups (WT-Veh, n = 10; WT-LPS, n =9; 5xFAD-Veh, n =9; 5xFAD-LPS, n = 8) in short-term memory test (STM) (Mean±SD, two-way ANOVA followed by Tukey’s test for multiple comparisons, P = 0.0269, *P < 0.05). C) Bar graph represents the preference of novel object in four different experimental groups (WT-Veh, n = 10; WT-LPS, n =9; 5xFAD-Veh, n =9; 5xFAD-LPS, n = 8) in long-term memory test (LTM) (Mean±SD, two-way ANOVA followed by Tukey’s test for multiple comparisons). D) Representative FACS plots for isolation of microglia using CD11b, CD45, and Cx3cr1 markers. E) Representative FACS plots for isolation of monocytes using Ly6C, CD11b, and CD115 markers.

### Novel object recognition task

The memory of the mice for the original object was assessed by comparing the number of times spent exploring the novel object with that spent exploring the familiar one. The cleaning procedure and the objects selected were similar to as previously described^11^. Before training, mice were allowed to explore the empty arena for 5 minutes to measure possible effects from the treatments on locomotion. Following this, two identical objects were placed in the center of the area for 15 minutes. Retention tests were then performed 1 h and 24 h after the training session to assess short-term memory and long-term memory, respectively. The mice were returned to the arena for a 10-minute session with the replacement of one original object with a novel one. The time spent exploring each object was recorded using Anymaze software (Stoelting Co, Wood Dale, IL), and the relative exploration of the novel object was calculated as a discrimination index [DI = (t_novel_-t_familiar_)/(t_novel_+t_familiar_)]. The criteria for exploration was based strictly on active exploration, defined as directing the nose toward the object at a distance of ±1.5cm and/or touching the object with the nose or vibrissae. Circling or sitting on the object were not considered exploratory behaviors. All trials were performed by an experimenter blind to the drug treatments and/or manipulations.

### Tissue processing and hippocampus homogenization

Mice were deeply anesthetized with 5% isoflurane in oxygen. They were then transcardially perfused with ice-cold phosphate-buffered saline (PBS). One hemisphere was removed and fixed in 4% PFA for 24 h, followed by PBS with 30% sucrose for 3 days. Coronal sections were sliced (30 μm thickness) using a freezing microtome (LeicaSM2000DR) and collected in cold cryoprotectant solution (phosphate buffer with 30% sucrose and 30% ethylene glycol) at − 20 °C. The other hemisphere excluding cerebellum was either snap frozen in dry ice and kept at − 80 °C for later or kept in cold PBS for isolation of microglia. Hippocampus tissue was lysed in Trizol (Sigma-Aldrich) reagent, and the protein was extracted. The concentration of total protein was measured with the Pierce™ BCA Protein Assay Kit (ThermoFisher Scientific). Equal amount of protein was loaded for every samples in the analysis of cytokines and Aβ.

### Single cell sorting

Brain suspension was prepared using an established method from Miltenyi Biotec using the Neural Tissue Dissociation Kit (130-092-628, Miltenyi Biotec). Bone marrow was collected from femur and tibia. Microglia and monocytes were then enriched by using CD11b microbeads (130-093-634, Miltenyi Biotec) and the Monocyte Isolation Kit (130-100-629, Miltenyi Biotec), respectively. Next, cells were sorted by a FACSArial III cytometer (BD Biosciences) with FACS Diva software (BD Biosciences). A viability stain, propidium iodide (PI) (421301, BioLegend, 1:1000), was used to exclude dead cells. Anti-CD11b-APC (Biolegend, 1:200), anti-CD45-PE (Biolegend, 1:200) and anti-CX3CR1-Alexa fluor® 488 (Biolegend, 1:200) were utilized to sort microglia. Monocytes were sorted using three surface markers: CD115-APC-Cy7 (135531, BioLegend, 1:200), CD11b-FITC (101217, BioLegend, 1:200) and Ly6C-PE (LS-C772930, Lifespan Bioscience; 1:200). Data was analyzed with FlowJo™ software version 10.4.0 (Becton, Dickinson and Company; 2019).

### Single cell sequencing

The sorted cells were prepared according to the 10x Genomics Next GEM Single cell 3′Reagent Kit v3.1. Approximately 20,000 cells from each sample were directly loaded onto an Illumina NovaSeq 6000. Manufacturer specifications were followed for generation of the barcoded libraries. 400 Mio reads were achieved for every sample.

### Analysis of single cell RNA-seq data

10x Genomics results were supplied by the facility as barcodes, features and mtx matrix files separately for each of the eight samples. Most downstream analyses were carried out using the R package Seurat 3.2.3^12^ in R 4.0.2 (Core Team, 2020). The filtered 10x data was estimated and corrected for background RNA contamination using SoupX 1.4.8^13^. Doublets were estimated using Scrublet^14^. Cell cycle phases were estimated using Seurat. Ribosomal genes (Rpl-) were removed from downstream analyses. Features that had fewer than or equal to 10 counts across all cells were discarded. Features with detection (1 or more reads) in fewer than 10 cells were discarded. Ribosomal genes (Rpl-) were discarded. Data normalisation (SCT) was carried out across all samples using the R package sctransform 0.3.2^15^. Number of RNA counts per cell, cell cycle S score, cell cycle G2M score and percentage of mitochondrial reads (Mt-) were regressed out during normalisation. 5000 variable features were retained.

For visualisation, the SCT data was integrated over type-condition (WT-Vehicle, WT-LPS, 5xFAD-Vehicle, 5xFAD-LPS) batches. PCA was run on the integrated data. UMAP was run on the first 40 principal components. Minimum distance and spread were set to 0.3 and 3 respectively. Louvain clustering was used to cluster the cells into 22 groups. Some of the very closely related groups which showed 100% bootstrap support using hierarchical clustering were merged back in, yielding 19 clusters. The MAST differential gene expression (DGE) test (McDavid et al., 2020) in Seurat was used to identify marker genes to distinguish each of the 19 clusters from each other. DGE was always performed on log normalised raw counts while correcting for covariates, including RNA counts per cell, cell cycle S score, cell cycle G2M score and percentage of mitochondrial reads. Minimum percentage expression across groups was set to 0.25, log-fold change threshold was set to 0.25, maximum cells per group was restricted to 100 and minimum cells per group was set to 50.

Cell type identification was based on a combination of known marker genes from literature and markers identified through MAST. Pairwise differential gene expression between groups of interest (clusters, samples etc.) was also carried out using MAST on log-normalised raw counts corrected for covariates. Minimum percentage expression across groups was set to 0.25, and both positive and negative fold-change were detected. Maximum cells per group was restricted to 500, and minimum cells per group was set to 10.

### Multiplex cytokine ELISA

The concentrations of different cytokines in serum and hippocampus homogenate were measured with the Mesoscale Discovery platform system (MSD, Rockville, USA). The MSD U-Plex Metabolic Group 1 (mouse) Kit (BDNF, GM-CSF, IFN-γ, IL-1β, TNF-α, MCP-1, MIP-1α, MIP-1β, MIP-2, MMP-9; K15317K) was selected to evaluate the cytokine levels. Three different forms of β-amyloid (Aβ38, Aβ40 and Aβ42) were quantified using the V-Plex Plus Aβ Peptide Panel 1 (6E10) Kit (K15200G-1, MSD, Rockville, USA) from hippocampal homogenate to evaluate the effect of the treatment on plaques. The plates were read by a QuickPlex Q120 according to the manufacturer’s instructions. The data was analyzed with the MSD Discovery Workbench software. In total, 6-7 serum samples for each condition and at least 3 independent biological replicates for brain homogenates were analyzed. However several of the samples were under the lowest detection limit and were excluded from the final statistical analysis.

### Quantification and statistical analysis

The differences between experimental groups were analyzed with an unpaired *t* test or two-or three-way ANOVA followed by Tukey’s test for multiple comparisons. P□<□0.05 was considered as statistically significant. All statistical analysis was done using the GraphPad Prism 7.0 software for macOS Catalina (GraphPad Software, San Diego, CA, USA). Data are presented as mean ± SD. A confidence interval of 95% was set as the significance level. The exact P values are given in the figure legends.

## Results

### Isolation and identification of microglia and monocytes

Increasing evidence has shown the heterogeneity of the microglia and monocyte population^16–18^. To understand how early-life stress affects the immune system in CNS and the periphery long-term, we isolated myeloid cells, including microglia, from brain and monocytes from bone marrow (Fig1.A) from 6-month-old mice subjected to i.p. injection of LPS (1mg/kg) at postnatal day 9. Moreover, in order to study the link between early inflammation and AD, we used a common AD transgenic mouse model, 5xFAD. Microglia were first enriched from brain tissue using CD11b magnetic beads, and monocytes were run through a negative selection of various antibodies targeting lymphocytes and granulocytes. Microglia were then FACS sorted with CD45, CD11b and Cx3cr1 as markers (Fig.1C), while monocytes were purified with CD115, Ly6C and CD11b surface markers (Fig.1D). Four independent biological replicates per group were collected and pooled as one sample for each condition. Monocytes and microglia were isolated from the same mice in each group. A total of 19,016 microglia and 20,653 monocytes from 32 mice underwent single-cell RNA sequencing. 31,053 features were available after computational analysis. The background RNA contamination was low in all of the samples. The clustering was estimated to 19 clusters for all cells (Fig2.A). Surprisingly, our analysis clearly indicated cell-type specific localization, as microglia resided primarily in the left continent (cluster 1 to 8) and monocytes at the right (cluster 9 to 19). This is consistent with our cluster dendrogram for microglia and monocyte populations as it can be divided into two main clusters from the different subpopulations (Fig.2B and Fig.3D).

**Figure 2.**
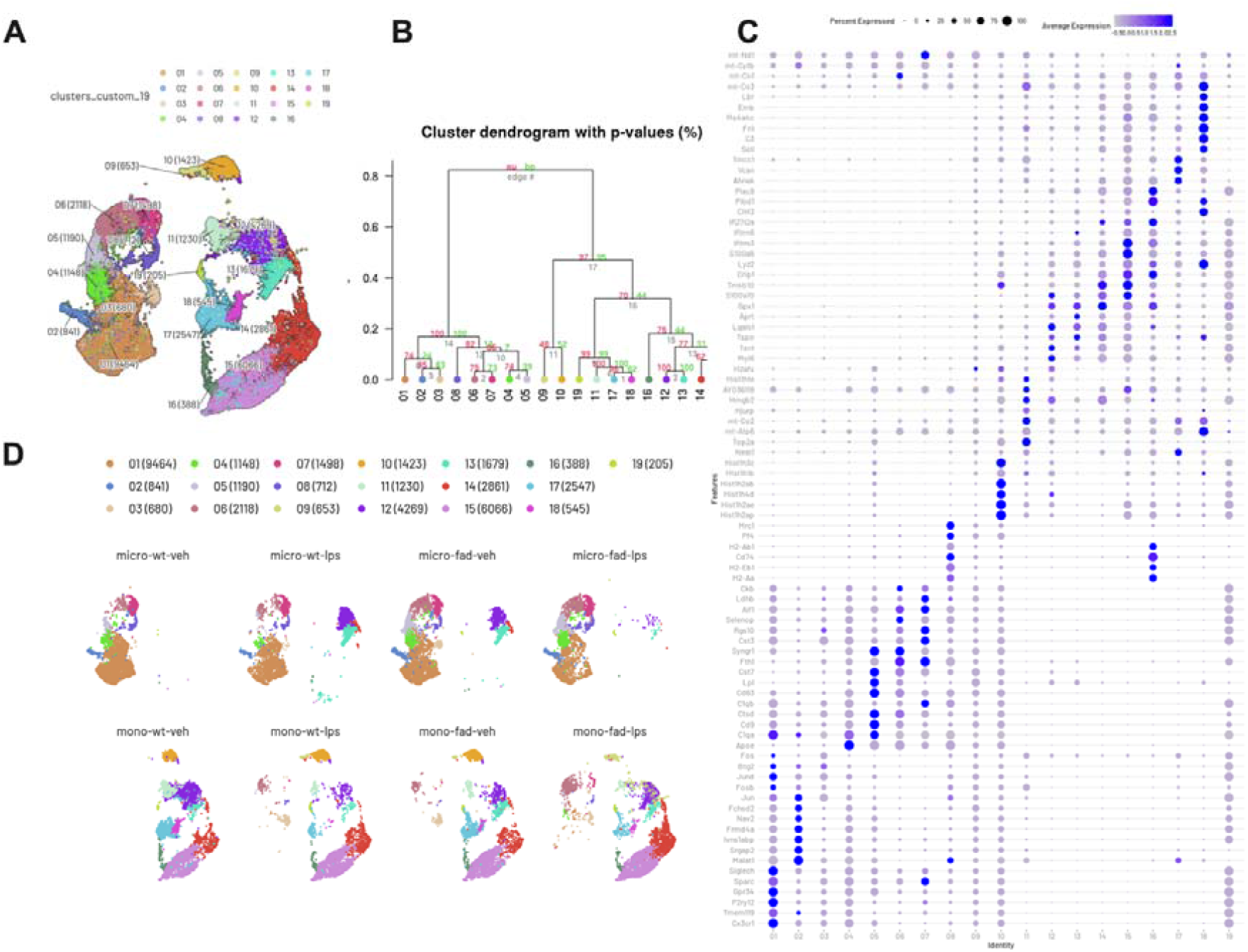
Single cell RNA-seq for 19 clusters found in all groups from microglia and monocytes. A) Clustering analysis of 39669 cells with 31053 features available from 4 mice per group for each cell type. B) Dendrogram shows the relation between different clusters. C) Dot plot shows the Top5 up-regulated genes for each cluster. D) Clustering analysis of microglia and monocytes from different experimental groups (n = 4 per group). micro: microglia, mono: monocyte.

### Specific markers identified in microglial clusters

Next, we looked at the dominant genes expressed at the highest levels for individual clusters to further investigate the functionality of various subsets. Cluster 1 was defined as homeostatic microglia due to highly expressed homeostatic genes, such as Cxcr1, Tmem119, P2ry12 and C1q^19,20^ (Fig.2C). The expression of Malat1 and Jun was pronounced in Cluster 2, which is a signature of inflammatory microglia mediated through the NF-κB signaling pathway^21,22^. From Clusters 4 to 6, the expression of Apoe was significantly increased and reached the peak in Cluster 5 (Fig.3A-C and Fig.5A). Many more AD risk genes were up-regulated in Cluster 5, including Trem2^23^, Tyrobp^24^ and Ctsd (Fig3.B). Several disease-associated microglia (DAM) markers CD9, Lpl, CD63^9^ were remarkably up-regulated in Cluster 5. Additionally, the expression of some homeostatic genes (P2ry12, Tmem119, Cx3cr1) were significantly reduced in Cluster 5 (Fig3.B). Therefore, we believe that we identified the DAM population in our data. A phenotype shift of microglia was found from Clusters 6 to 7 with down-regulation of Malat1 and Cx3cr1 (Fig3.C and E). The dot plot also indicated Cluster 8 highly expressed potential markers of perivascular macrophages, including Mrc1, H2-Ab and CD74^9,25^(Fig2.C).

**Figure 3.**
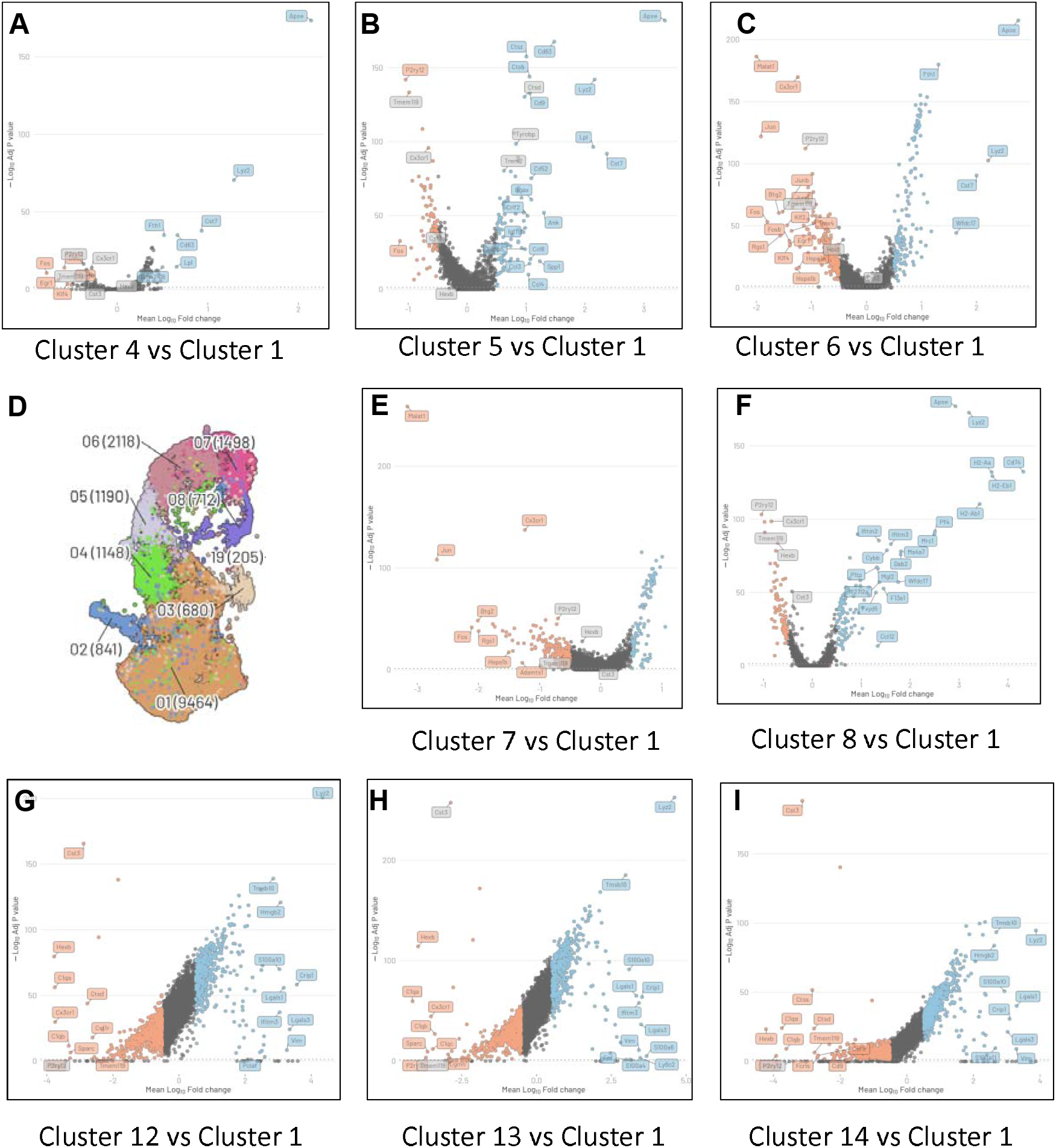
Volcano plots of comparisons between microglia clusters. A) Volcano plots shows the up- and down-regulated genes in Cluster 4 compared with Cluster 1. B) Volcano plots shows the up- and down-regulated genes in Cluster 5 compared with Cluster 1. C) Volcano plots shows the up- and down-regulated genes in Cluster 6 compared with Cluster 1. D) Clustering analysis on microglia for 9 clusters (n= 19016). E) Volcano plots shows the up- and down-regulated genes in Cluster 7 compared with Cluster 1. F) Volcano plots shows the up- and down-regulated genes in Cluster 8 compared with Cluster 1. G) Volcano plots shows the up- and down-regulated genes in Cluster 12 compared with Cluster 1. G) Volcano plots shows the up- and down-regulated genes in Cluster 13 compared with Cluster 1. G) Volcano plots shows the up- and down-regulated genes in Cluster 14 compared with Cluster 1. (Randomly selected 100 cells in each cluster for comparison.)

### Specific markers identified in monocyte clusters

We were also interested in potential markers for different monocyte clusters, which included clusters 10 to 19. Cluster 10 was enriched with Hist1h family genes, suggesting that chromatin remodeling and epigenetic alteration occurs within this subpopulation^26^. Two mitochondrial genes, mt-Co2 and mt-Atp6, were the distinctive genes for cluster 11 (Fig2.C). Levels of these two genes were previously found to be altered in blood from early-stage AD patients. However, this population is not unique to 5xFAD mice. Markers for clusters 12, 13 and a small part of 14 were found in both microglia and monocytes (Fig2.D). These three clusters were marked by Aprt and Lgals1(Fig2.C). Cluster 15 was identified by a large increase in S100a6 and Ifitm3 (Fig2.C), and Plbd1 and Plac8 were the top2 genes in Cluster 16 (Fig2.C). Vcan, Ahnak and Tmcc1 were genes upregulated in Cluster 17 (Fig2.C). The other three genes, Sell, fn1 and Ms4a4c, were described as the potential markers for Cluster 18. (Fig2.C).

### A distinct microglial population with a monocyte-like phenotype

We next investigated the small microglial island containing Clusters 12, 13 and 14 to understand why these populations localized closely with monocyte clusters. Strikingly, the small island was composed of only microglia from WT-LPS and 5xFAD-Vehicle groups (Fig2.D). A comparison was done between these clusters and Cluster 1 (homeostatic microglia) to find the differences. Lyz2 and Tmsb10 were highly up-regulated in these clusters as well as Lgals1 and Lgals3 (Fig3.G-I). These cells may have lost their microglia phenotype and adopted monocyte-like profiles because of a downregulation of microglial homeostatic genes (Hexb, Cst3, C1qa and Tmem119) (Fig3.G-I).

### Comparison between microglia and monocyte clusters

The similarity between microglia and monocytes has been an obstacle for researchers trying to distinguish them clearly. Additionally, setting up a boundary to identify these two cell types has been done arbitrarily. Therefore, we wondered how the transcriptomic profiles of common markers for microglia and monocytes appeared in our dataset. Five genes, including P2ry12, Cx3cr1, CD11b (Itgam), Fcrls and C1qa, were expressed more in microglia than in monocytes. In contrast, CD45 (Ptprc) and Ms4a family members, except Ms4a7, were at higher levels of expression in monocytes. Interestingly, several other genes were compared between clusters as well, and Lgals3 and lyz2 were upregulated more in all of the monocyte clusters (Figure 5A). CD11c (Itgax), a DAM marker, was particularly expressed in Cluster 5, which also had the highest expression of Apoe (Fig5.A). Cluster 8 distinctively expressed Ms4a7, the gene for a member of the transmembrane chemosensor family, which is involved in regulation of immune function and associated with Alzheimer’s disease risk. However, its role in the disease is not well known.

### Alteration of inflammatory cytokines in hippocampus and serum

To answer whether neuroinflammation occurred in the mice due to the AD pathology or early LPS treatment, we analyzed different inflammatory cytokines in hippocampus, a region closely linked to memory and cognition. GM-CSF was reduced significantly in mice with AD pathology (Fig4.A). However, MIP-1α was increased twofold in 5xFAD mice compared to WT mice (Fig4.C). Surprisingly, TNF-α was discovered in a genotype-dependent manner with significant down-regulation in 5xFAD mice (Fig4.D). However, it was remarkably increased in the serum after LPS treatment (Fig5.C). Moreover, IFN-γ was significantly reduced due to the LPS infection regardless of the genotype (Fig4.B). Strikingly, MIP-1β was dramatically low in the 5xFAD-Vehicle group but increased to a level similar to WT groups with early peripheral LPS injection. However, LPS infection did not alter the expression of MIP-1β in WT mice (Fig4.E). Another notable cytokine found in the serum was MMP-9, which was below the detection range in hippocampal samples. MMP-9 was found to be increased in all 5xFAD groups compared with WT (Fig5.B). These results suggest differing roles of inflammatory cytokines in the hippocampus compared to the periphery, especially TNF-α. We further looked into different forms of β-amyloid in hippocampus from 5xFAD-Vehicle and 5xFAD-LPS. No difference was detected for Aβ38 and Aβ40 (Fig4. F and G).

**Figure 4.**
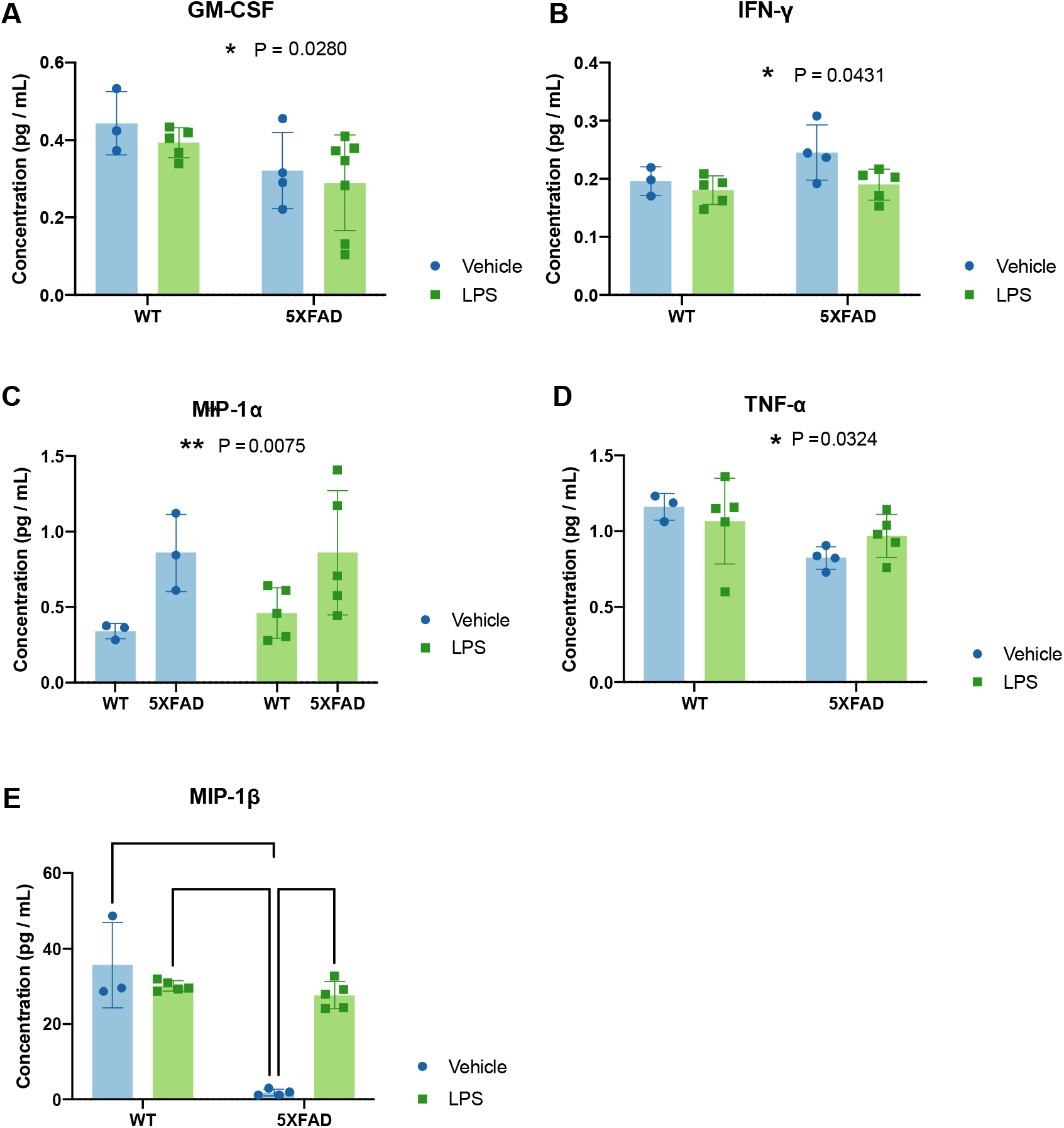
Cytokine measurement by multiplex ELISA in hippocampus area. A) Bar graph presents GM-CSF level in hippocampus from mice (WT, n=3; WT-LPS, n=5; 5xFAD, n=4; 5xFAD-LPS, n=5), (Two-way ANOVA followed by Tukey’s multiple comparison, *P < 0.05 due to genotype). B) Bar graph presents IFN-γ level in hippocampus from mice (WT, n=3; WT-LPS, n=5; 5xFAD, n=4; 5xFAD-LPS, n=5), (Two-way ANOVA followed by Tukey’s multiple comparison, *P < 0.05 due to LPS treatment). C) Bar graph presents MIP-1α level in hippocampus from mice (WT, n=3; WT-LPS, n=5; 5xFAD, n=4; 5xFAD-LPS, n=5), (Two-way ANOVA followed by Tukey’s multiple comparison, *P < 0.05 due to genotype). D) Bar graph presents TNF-α level in hippocampus from mice (WT, n=3; WT-LPS, n=5; 5xFAD, n=4; 5xFAD-LPS, n=5), (Two-way ANOVA followed by Tukey’s multiple comparison, *P < 0.05 due to genotype). E) Bar graph presents MIP-1β level in hippocampus from mice (WT, n=3; WT-LPS, n=5; 5xFAD, n=4; 5xFAD-LPS, n=5), (Two-way ANOVA followed by Tukey’s multiple comparison, ****P < 0.0001). F) Bar graph shows no difference at the level of β-amyloid 38 in hippocampus from mice (5xFAD, n=4; 5xFAD-LPS, n=5) (Mean±SD, Unpaired t-test). G) Bar graph shows no difference at the level of β-amyloid 40 in hippocampus from mice (5xFAD, n=4; 5xFAD-LPS, n=5) (Mean±SD, Unpaired t-test).

**Figure 5.**
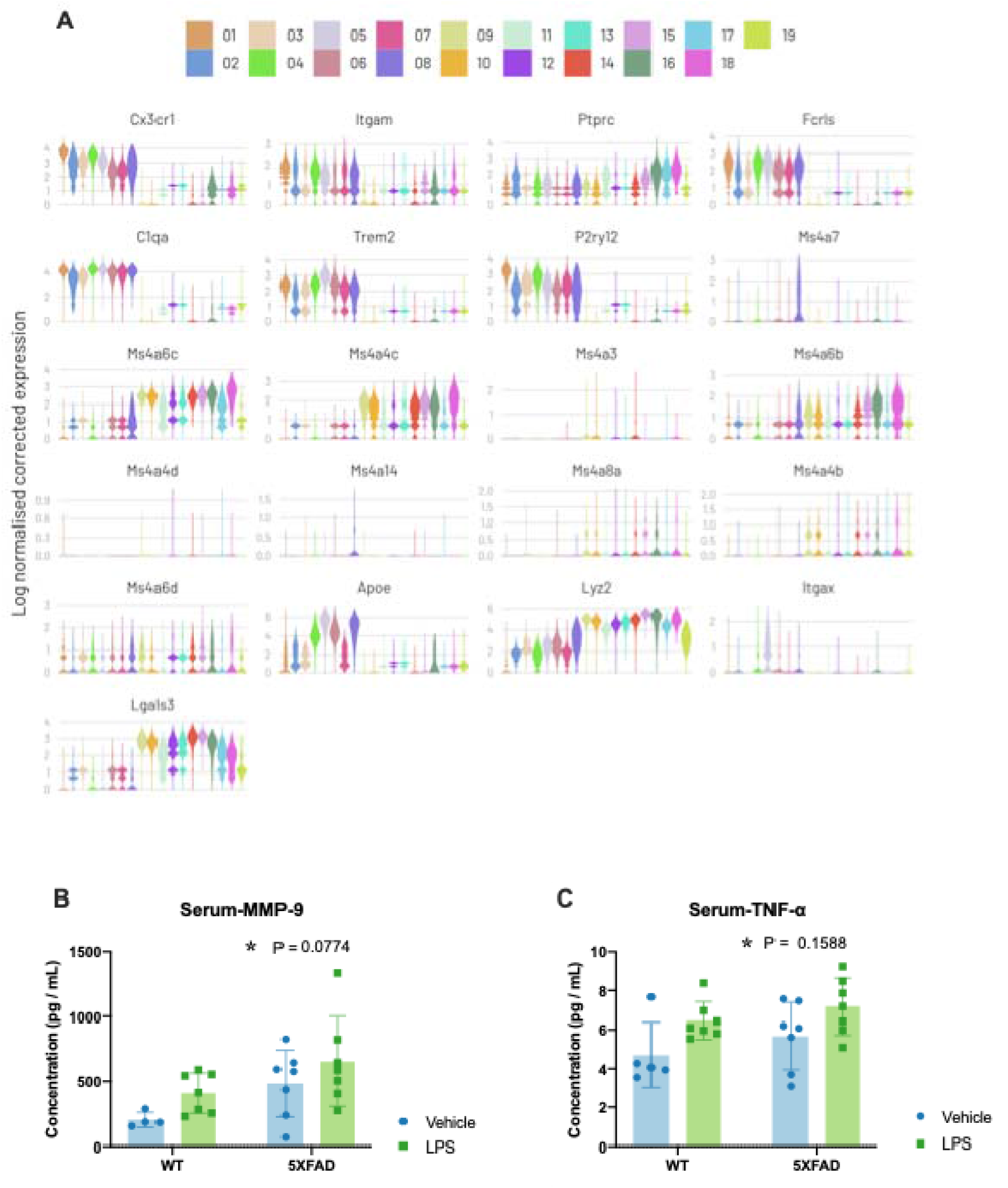
Expression level of common genes in microglia and monocyte. A) Violin plots presents homeostatic genes in microglia including Cx3cr1, P2ry12, C1qa and Fcrls as well as Alzheimer’s disease related genes such as Trem2, Apoe, Ms4a family, Lgals3 and DAM markers (Itgax). Common markers including Ptprc and Itgam. B) Bar graph presents MMP-9 level in serum samples (WT, n=4; WT-LPS, n=7; 5xFAD, n=7; 5xFAD-LPS, n=7), (Two-way ANOVA followed by Tukey’s multiple comparison, *P < 0.05 due to genotype and *P < 0.05 in WT-Vehicle vs 5xFAD-LPS). C) Bar graph presents TNF-α level in serum samples (WT, n=5; WT-LPS, n=7; 5xFAD, n=7; 5xFAD-LPS, n=7), (Two-way ANOVA followed by Tukey’s multiple comparison, *P < 0.05 due to genotype and *P < 0.05 in WT-Vehicle vs 5xFAD-LPS).

### LPS does not affect cognition in WT and 5xFAD mice

In order to investigate how early infection modulates AD pathology long-term, we performed i.p. injection of LPS (1mg/kg) at postnatal day 9 in WT and 5xFAD mice. There was no increase in mortality after one dose of LPS. We sought to test the cognitive impairment in these mice, as synapse loss is one of the hallmarks in AD and contributes to memory impairment in different aspects. A novel object recognition task (see Fig1.A for the experiment outline) was conducted at 6 months of age to evaluate the long-term effect of peripheral inflammation on retention capacity of 5xFAD and WT mice. The retention intervals were defined as 1 h as a short-term memory (STM) test and 24 h as a long-term memory (LTM) test. No significant difference was found between genotype or treatment using three-way ANOVA (Fig1B). However, in vehicle groups, we saw a significant decrease in preference to the novel object in 5xFAD mice (Fig.1B). Interestingly, early infection in the periphery disrupted this phenomenon. This evidence highlights the cognitive alteration in mice due to LPS stress early in life.

## Discussion

The immune response and related pathways are frequently elicited in AD and other neurodegenerative diseases. With an increasing number of researchers using single cell RNA-seq as a means to study this question, the heterogeneity of microglia and monocytes was uncovered and could be defined with diverse markers with potential roles in different signaling cascades^8,9,16,17^. The neuroinflammation in AD has been associated with cytotoxicity, and microglia have been implicated to play an important role during the progression of disease^27–29^. In contrast, some studies have attributed a positive role to infiltrating monocytes as they may aid in the clearance of toxins and β-amyloid from the brain^9,30,31^. Moreover, the connection between monocytes in peripheral and microglia in CNS is not yet clear.

In this study, we compared the immune response of monocytes and microglia in mice that experienced an acute infection systematically as well as developed AD pathology as a chronic inflammation in the brain. Our results suggest these two conditions triggered different immune responses in microglia and monocytes. Here, we identify a novel subtype of microglia with a monocyte-like transcriptomic profile and decipher their reaction to different inflammatory stimuli. This distinct subtype exists in both WT mice after early-life systemic infection and 5xFAD without early-life infection. In other words, the conditions when only one stimulus challenged the immune system. Further investigation of this subtype revealed potential markers, including Lyz2, S100a6, Lgals3 (galectin-3), and Lgals1 (galectin-1), as labels that could be used for immunostaining or qPCR to further validate and examine the functionality of this subpopulation in the brain. Lyz2 is primarily involved in the clearance of bacteria, and S100a6 may lead to apoptosis through activation of the RAGE signaling pathway^32^. A previous study showed that galectin-1 could be a protective marker for bone-marrow derived macrophages^33^. In addition, galetin-3 has been regarded as an activated microglia marker and implicated as a ligand for Trem2 with up-regulation corresponding to AD progression^34^. Altogether, these potential markers suggest an active role of microglia in response to inflammation.

In total, we identified 19 clusters of monocytes and microglia. Among them, homeostatic microglia were contained in Cluster 1, and DAM were found in in Cluster 5. Perivascular macrophages were localized in Cluster 8. Besides DAM genes, Apoe and CD11c were expressed the most in Cluster 5. DAM are localized in the vicinity of plaques and phagocytose β-amyloid in CNS^9^. These findings highlight the protective role of DAM in AD. Interestingly, this cluster appears primarily in 5xFAD mice with some DAM in WT as well but very few appeared in the WT-LPS group (Fig2.D). This implies that DAM may be more responsive to events requiring phagocytosis, such as, plaques and may not respond effectively to LPS stimulation. Further comparisons of monocytes are needed to investigate the different responses in the periphery.

To examine the inflammation in the CNS and periphery, different cytokines were measured to determine the effects of early-life stress and chronic inflammation from AD pathology. GM-CSF can induce programmed cell death in brain tissue and has been previously shown to be increased in cerebrospinal fluid and serum from AD and vascular dementia patients^35^. These findings go against what we found from our hippocampal samples. This different suggests that GM-CSF has region specificity or may play different roles in mice compared to humans. MIP-1α has been found to be increased in vessels from AD patients, and here we confirmed the same effect in hippocampus^36^. Strikingly, MIP-1β (CCL4) was rescued by the systemic LPS infection in 5xFAD mice. CCL4 has been demonstrated to be closely connected with ageing ^18^, but its role in neurodegenerative diseases has not been thoroughly investigated. In our data analysis, CCL4 was highly upregulated in Cluster 5, where DAM localized, suggesting a potential neuroprotective role in microglia. Increasing the number of mice in the groups and immunostaining of the brain sections could help to further validate these results. Another interesting cytokine that came up in our analyses was TNF-α due to differences in its secretion in CNS compared to the periphery. TNF-α is a pro-inflammatory cytokine and an essential component in neuroinflammation that has been associated with several neurological diseases. However, we saw a long-lasting effect in the serum after LPS infection but not in hippocampus, which suggests differing roles of TNF-α in the CNS versus periphery. One study showed that the transmembrane form of TNF-α may have a neuroprotective role, while the soluble form may be involved in promoting inflammation^37^. Therefore, different end effects may arise depending on the result of the competition between these two forms. In our hippocampal samples, we have homogenated the tissue, so both forms of TNF-α are likely represented by the measurement. However, in the serum, we believe the soluble form of TNF-α is the predominant form.

We further investigated the effect of early-life stress on memory formation. Behavioral tests were performed to look at STM and LTM. Indeed, inflammation that occurred in the early stages of brain development disrupted the cognition decline in 5xFAD mice. Few studies have looked at the long-term effect of stress in cognition. However, our work is limited as more mice are needed per group for the tests as there are large variations within the groups. However, measurements of the toxic forms of β-amyloid 38 and 40 showed no difference between 5xFAD and 5xFAD-LPS mice. The other toxic form, β-amyloid 42, will require further analysis, including immunostaining of plaques and microglia to evaluate the pathology and neuroinflammation in detail.

Taken together, our results demonstrate a subpopulation of microglia that adopt a monocyte-like profile, as defined by the expression of Lyz2, Tmsb10, Lgals1and Lgals3. This unique subset appeared in response to early systemic LPS infection and AD pathology but diminished in the presence of double neuroinflammatory stimuli. DAM and perivascular macrophages were successfully mapped in our clustering. Different cytokines were altered in the brain and periphery. GM-CSF, MIP-1α and TNF-α were changed in an AD-dependent manner in hippocampus. MIP-1β and IFN-γ were changed upon early LPS stimulation. In the periphery, MMP-9 was increased significantly in 5xFAD mice, and TNF-α was elevated after LPS treatment in both WT and transgenic mice. We have observed long-lasting effects of early-life stress, including activation of inflammation in the periphery as well as cognitive alterations. Future work would include validation of the results in AD and sepsis patients.

## Declarations

### Authors’ contributions

YY designed the studies, performed experiments, and wrote the manuscript. TD participated in data analysis, interpreted the results and supported the study with funding. HZ did computational analysis of single-cell RNA-seq data. SB helped with data interpretation, designed and analyzed the cognitive behavioral tests and performed it with YY. MGC, BBK, XW did immunostaining, confocal microscopy and data quantification. LCF contributed to quantification of Aβ pathology. RFC performed qPCR and analyzed the results. All authors have read and approved the final manuscript.

## Acknowledgements

We acknowledge technical support for animal handling from Bodil Israelsson, flow cytometry from Anna Hammarberg at Multipark Lund University, Mouse Behavioral Platform at Multiplark Lund University, Lund Univeristy Bioimaging Center.

## References

1. Bechade, C., Cantaut-Belarif, Y. & Bessis, A. Microglial control of neuronal activity. Front Cell Neurosci (2013) doi:10.3389/fncel.2013.00032.

2. Labzin, L. I., Heneka, M. T. & Latz, E. Innate Immunity and Neurodegeneration. Annu Rev Med (2018) doi:10.1146/annurev-med-050715-104343.

3. Khoury, J. E. et al. Ccr2 deficiency impairs microglial accumulation and accelerates progression of Alzheimer-like disease. Nat Med 13, 432–438 (2007).

4. Gordon, S. & Taylor, P. R. Monocyte and macrophage heterogeneity. Nat Rev Immunol 5, 953–964 (2005).

5. Reed-Geaghan, E. G., Croxford, A. L., Becher, B. & Landreth, G. E. Plaque-associated myeloid cells derive from resident microglia in an Alzheimer’s disease model. J Exp Medicine 217, (2020).

6. Rantakallio, P., Jones, P., Moring, J. & Wendt, L. V. Association between central nervous system infections during childhood and adult onset schizophrenia and other psychoses: A 28-year follow-up. Int J Epidemiol (1997) doi:10.1093/ije/26.4.837.

7. Frost, P. S. et al. Neonatal infection leads to increased susceptibility to A$\beta$ oligomer-induced brain inflammation, synapse loss and cognitive impairment in mice. Cell Death Dis (2019) doi:10.1038/s41419-019-1529-x.

8. Reyes, M. et al. An immune-cell signature of bacterial sepsis. Nat Med 26, 333–340 (2020).

9. Keren-Shaul, H. et al. A Unique Microglia Type Associated with Restricting Development of Alzheimer’s Disease. Cell 169, 1276-1290.e17 (2017).

10. Varol, D. et al. Dicer Deficiency Differentially Impacts Microglia of the Developing and Adult Brain. Immunity 46, 1030-1044.e8 (2017).

11. Bachiller, S., del-Pozo-Martín, Y. & Carrión, Á.M. L1 retrotransposition alters the hippocampal genomic landscape enabling memory formation. Brain Behav Immun 64, 65–70 (2017).

12. Stuart, T. et al. Comprehensive Integration of Single-Cell Data. Cell 177, 1888-1902.e21 (2019).

13. Young, M. D. & Behjati, S. SoupX removes ambient RNA contamination from droplet-based single-cell RNA sequencing data. Gigascience 9, giaa151.(2020).

14. Wolock, S. L., Lopez, R. & Klein, A. M. Scrublet: Computational Identification of Cell Doublets in Single-Cell Transcriptomic Data. Cell Syst 8, 281-291.e9 (2019).

15. Hafemeister, C. & Satija, R. Normalization and variance stabilization of single-cell RNA-seq data using regularized negative binomial regression. Genome Biol 20, 296 (2019).

16. Grubman, A. et al. A single-cell atlas of entorhinal cortex from individuals with Alzheimer’s disease reveals cell-type-specific gene expression regulation. Nat Neurosci 22, 2087–2097 (2019).

17. Friedman, B. A. et al. Diverse Brain Myeloid Expression Profiles Reveal Distinct Microglial Activation States and Aspects of Alzheimer’s Disease Not Evident in Mouse Models. Cell Reports 22, 832--847 (2018).

18. Hammond, T. R. et al. Single-Cell RNA Sequencing of Microglia throughout the Mouse Lifespan and in the Injured Brain Reveals Complex Cell-State Changes. Immunity 50, 253-271.e6 (2019).

19. Merino, J. J., Muñetón-Gómez, V., Alvárez, M.-I. & Toledano-Díaz, A. Effects of CX3CR1 and Fractalkine Chemokines in Amyloid Beta Clearance and p-Tau Accumulation in Alzheimer,s Disease (AD) Rodent Models: Is Fractalkine a Systemic Biomarker for AD? Curr Alzheimer Res 13, 403–412 (2016).

20. Butovsky, O. et al. Targeting miR[155 restores abnormal microglia and attenuates disease in SOD1 mice. Ann Neurol 77, 75–99 (2015).

21. Waetzig, V. et al. c[Jun N[terminal kinases (JNKs) mediate pro[inflammatory actions of microglia. Glia 50, 235–246 (2005).

22. Zhou, H.-J. et al. Long noncoding RNA MALAT1 contributes to inflammatory response of microglia following spinal cord injury via the modulation of a miR-199b/IKKβ/NF-κB signaling pathway. Am J Physiol-cell Ph 315, C52–C61 (2018).

23. Singaraja, R. TREM2: a new risk factor for Alzheimer’s disease. Clin Genet 83, 525–526 (2013).

24. Pottier, C. et al. TYROBP genetic variants in early-onset Alzheimer’s disease. Neurobiol Aging 48, 222.e9-222.e15 (2016).

25. Li, Q. & Barres, B. A. Microglia and macrophages in brain homeostasis and disease. Nat Rev Immunol 18, 225–242 (2018).

26. Laan, L. et al. DNA methylation changes in Down syndrome derived neural iPSCs uncover co-dysregulation of ZNF and HOX3 families of transcription factors. Clin Epigenetics 12, 9 (2020).

27. Mosher, K. I. & Wyss-Coray, T. Microglial dysfunction in brain aging and Alzheimer’s disease. Biochem Pharmacol 88, 594–604 (2014).

28. Yuan, P. et al. TREM2 Haplodeficiency in Mice and Humans Impairs the Microglia Barrier Function Leading to Decreased Amyloid Compaction and Severe Axonal Dystrophy. Neuron 90, 724–739 (2016).

29. Prinz, M. & Priller, J. Microglia and brain macrophages in the molecular age: from origin to neuropsychiatric disease. Nat Rev Neurosci 15, 300–312 (2014).

30. Jay, T. R. et al. TREM2 deficiency eliminates TREM2+ inflammatory macrophages and ameliorates pathology in Alzheimer’s disease mouse modelsTREM2 expression and function in AD. J Exp Medicine 212, 287–295 (2015).

31. Koronyo, Y. et al. Therapeutic effects of glatiramer acetate and grafted CD115+ monocytes in a mouse model of Alzheimer’s disease. Brain 138, 2399–2422 (2015).

32. Xia, C., Braunstein, Z., Toomey, A. C., Zhong, J. & Rao, X. S100 Proteins As an Important Regulator of Macrophage Inflammation. Front Immunol 8, 1908 (2018).

33. Krautter, F. et al. Characterisation of endogenous Galectin-1 and -9 expression in monocyte and macrophage subsets under resting and inflammatory conditions. Biomed Pharmacother 130, 110595 (2020).

34. Boza-Serrano, A. et al. Galectin-3, A Novel Endogenous Trem2 Ligand, Regulates Inflammatory Response and Aβ Fibrilation in Alzheimer’s Disease. Biorxiv 477927 (2018) doi:10.1101/477927.35.

35. j.1600-0404.2001.103003166.x.pdf. undefined.

36. Tripathy, D., Thirumangalakudi, L. & Grammas, P. Expression of Macrophage Inflammatory Protein 1-α is Elevated in Alzheimer’s Vessels and is Regulated by Oxidative Stress. J Alzheimer’s Dis 11, 447–455 (2007).

37. Taoufik, E. et al. Transmembrane tumour necrosis factor is neuroprotective and regulates experimental autoimmune encephalomyelitis via neuronal nuclear factor-κB. Brain 134, 2722–2735 (2011).

